# Sex-specific electrophysiology and cholinergic responses underlie differential mechanisms of arrhythmia vulnerability in rabbit atria

**DOI:** 10.64898/2026.02.18.706701

**Authors:** Charlotte E. R. Smith, Lianguo Wang, Amanda M. Guevara, Lilian R. Mott, Haibo Ni, Eleonora Grandi, Crystal M. Ripplinger

## Abstract

**Background:** Sex differences in the epidemiology of atrial fibrillation are well-documented; however, the underlying mechanisms remain poorly understood. This gap in knowledge is compounded by limited data on sex-specific atrial electrophysiology in the absence of disease.

**Objectives:** The aim of this study was to investigate sex differences in atrial electrophysiology and arrhythmia susceptibility in a translationally-relevant rabbit model.

**Methods:** Dual optical mapping of transmembrane voltage and Ca^2+^ was performed on intact atria of young (3.5-5 months) male and female rabbit hearts. Baseline atrial electrophysiology and arrhythmia susceptibility were investigated using rapid pacing and premature stimulation and further tested with the parasympathomimetic carbachol. Sex and regional differences in gene expression were assessed using qPCR.

**Results:** Females exhibited similar action potential duration (APD), but greater APD heterogeneity across the atria at slower rates, along with longer Ca^2+^ transient durations compared to males. Greater APD heterogeneity in females was rate-dependent and comparable to males at faster pacing frequencies; however, it was associated with greater susceptibility to transient reentrant arrhythmias with premature stimuli. After carbachol application, males had heightened vulnerability to arrhythmia. This was associated with cholinergic-mediated APD shortening in both atria in males, but only in the right atrium in females. Sex differences in carbachol responses were linked to variations in muscarinic receptor and acetylcholine-activated potassium channel gene expression.

**Conclusions:** These findings demonstrate sex and regional differences in atrial electrophysiology at baseline and in response to cholinergic stimulation in the healthy heart that may contribute to sex-specific mechanisms of arrhythmia.

## INTRODUCTION

Atrial fibrillation (AF), the most common cardiac arrhythmia, is characterized by rapid and irregular electrical activity that reduces atrial function and causes dyssynchronous blood pumping. Consequently, AF not only causes significant morbidity but is also associated with greater risk of mortality through heart failure and thromboembolic events such as stroke and myocardial infarction. Like many cardiovascular diseases, AF exhibits significant sex differences in onset, symptoms and treatment outcomes.^1–3^ While the overall incidence is similar between sexes, the age of onset in men is ∼8 years earlier than women, yet women experience more debilitating symptoms, less successful outcomes from ablation and drug therapies, and face a higher risk of adverse events and mortality.^1–7^ Despite these clinical disparities, the mechanisms underlying sex differences in AF remain poorly understood. In particular, due to the historical underrepresentation of females in both preclinical and clinical research, little is known about sex differences in the healthy atria and how these may predispose individuals to the sex-specific manifestations and outcomes observed in the disease state.

Recent studies have begun to characterize sex-specific differences in baseline atrial electrophysiology using isolated myocytes and tissue preparations, identifying variations in action potential (AP) morphology and Ca^2+^ cycling that may underly differences in arrhythmia susceptibility.^3,8–17^ While these investigations have provided valuable insights at the cellular and tissue levels, many do not consider rate-dependent properties, interatrial heterogeneity, or utilize the intact atria, factors that are essential for understanding dynamic electrophysiological behavior and elucidating complex arrhythmia mechanisms.

To address this gap and bridge the divide between cellular studies and translational electrophysiological outcomes, we investigated sex differences in atrial electrophysiology and arrhythmia susceptibility in the intact rabbit heart, which has similar electrophysiological characteristics to humans.^15,18^ Using dual optical mapping of transmembrane voltage and intracellular Ca^2+^, we found that young female atria exhibited similar AP durations (APD), but longer Ca^2+^ transient durations (CaTD) compared to males. Although mean APD was comparable between groups, female atria showed greater APD heterogeneity at slower pacing rates, primarily due to prolonged APD in the right atrium (RA) and interatrial region (IAR). This sex difference in heterogeneity was rate-dependent and diminished at faster pacing rates; however, it was associated with greater susceptibility to transient reentrant arrhythmias when using premature stimulation (S1-S2) pacing protocols. We also assessed vulnerability to arrhythmia following application of the parasympathomimetic carbachol and observed tendency for elevated arrhythmia risk under cholinergic stimulation in males. This was associated with greater carbachol-induced APD shortening across the atria in males compared to females and was linked with sex and chamber differences in muscarinic receptor and acetylcholine-activated potassium channel gene expression. Together, these data reveal sex and regional differences in atrial electrophysiology both at baseline and in response to cholinergic activation, which may contribute to sex-specific mechanisms underlying atrial arrhythmogenesis.

## METHODS

### Ethical approval

All procedures were approved by the Animal Care and Use Committee of the University of California, Davis and were conducted according to the Guide for the Care and Use of Laboratory Animals published by the National Institutes of Health (NIH publication no. 85-23, revised 2011).

### Heart extraction and perfusion

Young adult male and female New Zealand White rabbits (3.5 – 5 months old, 2.8 – 3.4 kg) were purchased from Charles River and housed individually for at least two weeks prior to use, with *ad libitum* access to food and water. Rabbit hearts were isolated as previously described.^19–22^ Briefly, animals were heparinized (1000 IU) and then euthanized by overdose of pentobarbital sodium (>100 mg/kg) via a catheter placed in the marginal ear vein. Hearts were extracted and submerged in ice-cold cardioplegia solution (containing in mM: 110 NaCl, 16 KCl, 16 MgCl_2_, 10 NaHCO_3_ and 1.2 CaCl_2_) prior to aortic cannulation and perfusion at 37°C with Tyrode’s solution (containing in mM: 128.2 NaCl, 20 NaHCO_3_, 11.1 glucose, 4.7 KCl, 1.3 CaCl_2_, 1.19 NaH_2_PO4 and 1.05 MgCl_2_). Hearts were perfused posterior side up to permit optimal atrial imaging and the excitation-contraction uncoupler blebbistatin (20 µM, Tocris Bioscience) was added to the perfusate to eliminate motion artifacts during optical recordings.^23^ Excess fat and vascular tissue were trimmed from around the IAR, and the atria were carefully pinned out (Fig. 1A). Three Ag/AgCl needle electrodes were placed in the dish for continuous *ex vivo* ECG recording via Labchart (ADInstruments). Perfusion pressure was maintained at 60 – 80 mmHg by adjusting perfusion flow rate (∼30 mL/min).

**Figure 1.**
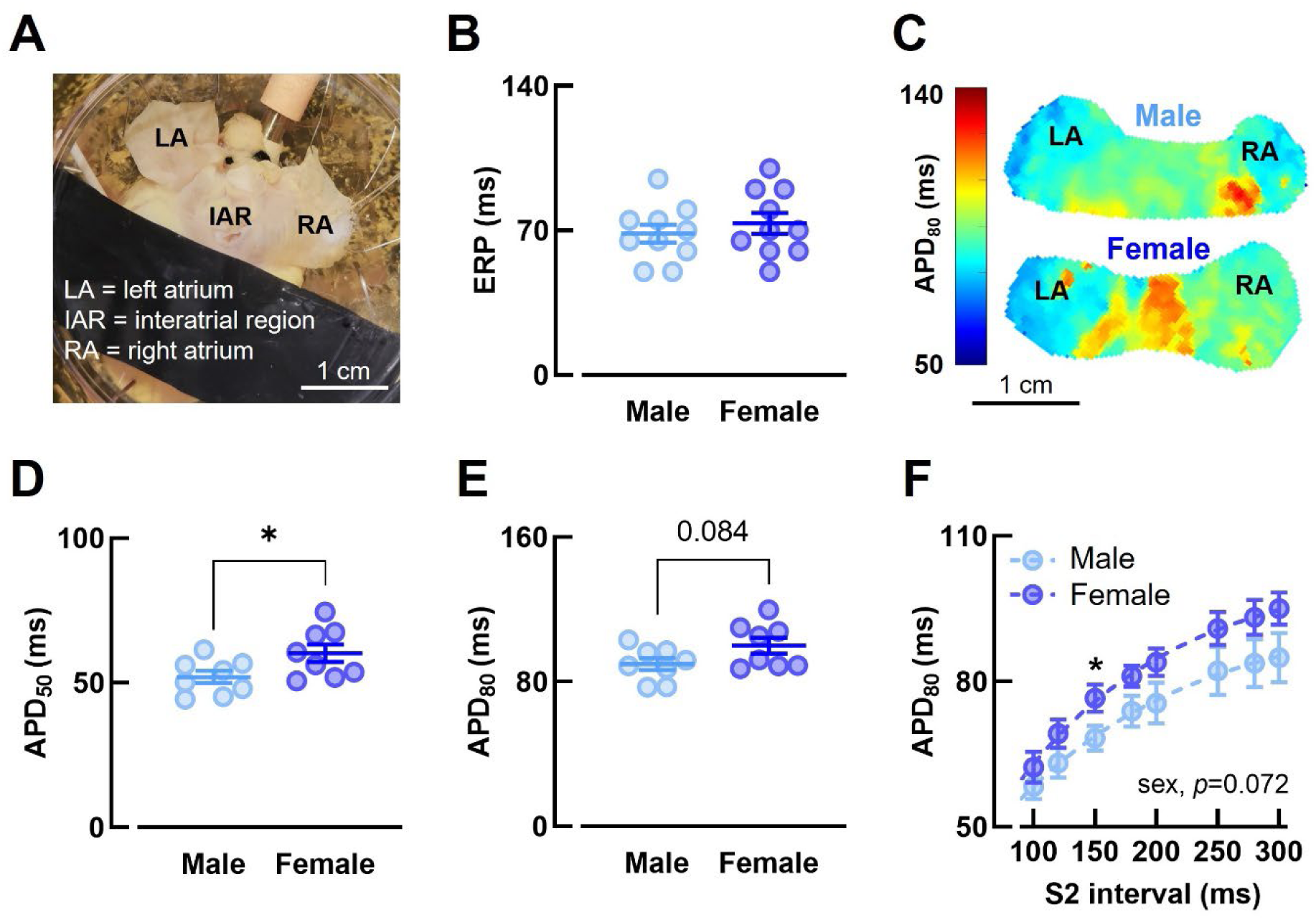
Atrial electrophysiological parameters are similar in male and female rabbits. **(A)** Example atrial mapping preparation. **(B)** ERP, **(C)** example APD_80_ maps, **(D)** APD_50_, **(E)** APD_80_ and **(F)** APD_80_ restitution in male and female rabbit atria. B, N = 10/group. C, D, E and F, N=8/group. Statistical significance assessed using unpaired *t*-test (B, D and E) and two-way mixed effects model (F). B and F, S1 interval = 280 ms. C, D and E, BCL = 300 ms. APD_50_ = action potential duration at 50% repolarization, APD_80_ = action potential duration at 80% repolarization, BCL = basic cycle length, ERP = effective refractory period.

### Dual optical mapping

Dual optical mapping was performed as described previously.^19,20,24–28^ Hearts were loaded with the Ca^2+^ indicator Rhod-2 AM (Biotium), 1 mg/mL in 0.5 mL DMSO containing 10% pluronic acid, followed by the voltage-sensitive dye RH237 (Biotium), 50 µL 1 mg/mL in DMSO, via the perfusion system. The atrial epicardial surface was excited with two LED light sources filtered at 531 ± 20 nm (LEX-2, SciMedia). Emitted fluorescence was captured with a THT-macroscope (SciMedia) and split by a dichroic mirror at 630 nm (Omega Optical). Voltage signals were filtered using a longpass filter at 700 nm and Ca^2+^ signals were filtered using a 590 ± 16 nm bandpass filter. Signals were recorded with two CMOS cameras (MiCam Ultima-L, SciMedia) at a sampling rate of 1 kHz, 100 x 100 pixel resolution and field of view of 3.1 x 3.1 cm, providing a pixel size of 310 µm. To minimize ventricular signal interference, the ventricles were covered below the coronary sulcus with photoblocking material for the duration of the experiment (Figure 1A). Pacing was performed from a bipolar pacing electrode placed on the RA epicardium. Baseline Ca^2+^ and voltage recordings were obtained during pacing at a basic cycle length (BCL) of 300 ms. Continuous pacing was performed to assess rate-dependence of heterogeneity and alternans susceptibility and S1-S2 pacing protocols were performed to assess restitution and effective refractory period (ERP).^29^ Pacing started from a BCL of 300 ms or S1 interval of 280 ms for 12 beats then the BCL or S2 interval was progressively decreased from 300 to 250, 200, and 180 ms, followed by further 10 ms decrements until loss of capture. Susceptibility to arrhythmia was assessed using burst pacing with pause (BCL = 100 and 50 ms) and the premature stimulation (S1-S2) protocols detailed above. All protocols were also repeated in the presence of the parasympathomimetic carbachol (0.5 µM, Sigma-Aldrich). Carbachol was added to the perfusate and allowed to circulate for 10 minutes prior to resuming data recording and repeating the pacing protocols. Where sustained arrhythmias occurred, termination was first attempted using overdrive pacing at shorter BCLs, followed, if necessary, by defibrillation at 2 V using a Heartstart XL defibrillator (Philips).

### Optical mapping data analysis

Optical data were analyzed using Electromap^30^ and Optiq (Cairn Research) as previously described.^24,31,32^ APD and CaTD were calculated at 50 and 80% repolarization/decay minus activation time (with activation time calculated at 50% of peak amplitude). Measurements were taken either across the entire atria or from a 9×9 pixel region of interest in the left atria (LA), RA or IAR as specified. These same 9×9 regions of interest were used to generate representative traces. Heterogeneity was assessed by calculating the standard deviation of APD and CaTD across the entire atria. Restitution curves were generated by plotting APD₈₀ against the corresponding S2 interval. Restitution steepness was assessed by fitting a linear regression to the data within the 200–100 ms S2 interval range and comparing the resulting slopes between groups. The emergence of Ca^2+^ transient (CaT) and AP alternans was determined using spectral methods as previously described, with the threshold for alternans defined as the BCL at which significant alternans emerged (minimum spectral magnitude ≥ 2).^21,22,33^ ERP was defined as the longest S2 interval that failed to elicit a response. The relative recovery of Ca²⁺ release was calculated during premature stimulation protocols where the CaT amplitude during the S2 beat was normalized to the amplitude of the S1 CaT.^33^ Heart rate was analyzed using the continuous ECG recordings obtained during mapping using shape recognition algorithms in LabChart (ADInstruments). Arrhythmias were categorized as ectopic or reentrant, and as transient or sustained based on optical and ECG recordings. Classification was determined by their mode of onset, duration and termination characteristics whereby transient arrhythmias self-terminated spontaneously and sustained arrhythmias required pacing overdrive or defibrillation for termination.

### Gene expression

Upon completion of mapping experiments, tissue was harvested from the LA and RA, flash-frozen in liquid nitrogen and stored at −80°C. To assess the expression of key ion channel, receptor and Ca^2+^-handling genes, qPCR was performed by the University of California Davis School of Veterinary Medicine Real-time PCR Research and Diagnostics Core Facility. Total nucleic acid (DNA and RNA) was extracted from approximately 10 mg of tissue using the QIAamp 96 DNA QIAcube HT Kit (QIAGEN) on a Qiagen QIAcube HT semi-automated, filter-based extraction system. cDNA synthesis was performed using SuperScript IV Vilo Mastermix with EZDNase (Thermo Fisher Scientific) using 10 µL of extracted total nucleic acid as the template. To confirm successful DNase digestion, a subset of samples was assayed for eukaryotic 18S post DNase treatment but prior to cDNA synthesis. All samples showed sufficient DNA elimination. qPCR was performed using custom-designed TaqMan Array Plates and TaqMan Gene Expression Master Mix (Thermo Fisher Scientific) according to the TaqMan Gene Expression Assays – Taqman Array Plates User Guide for 384-well Specialty TaqMan Array Plates. Reactions were run on a QuantStudio 7 Pro thermocycler (Applied Biosystems) using the following cycling conditions: 2 min at 50°C, 10 min at 95°C, 40 cycles of 95°C for 15 sec and 60°C for 1 min. Quantification cycle (Ct) values were extracted during the annealing phase using a baseline of 3-15 and a threshold of 30,000 for all genes, except for *KCNJ4* (inward rectifier K^+^ channel subunit Kir2.3), for which a threshold of 20,000 was applied. Each array included 16 genes of interest and three reference genes (Table 1). Of the 16 target genes, all showed successful amplification except *KCNJ4* which was below the threshold for reliable detection. All reactions were performed in triplicate, and relative gene expression was calculated using the 2^-ΔΔ*C*^_t_ method, normalized to the geometric mean of the reference genes and to expression levels in the male LA.^34^

**Table 1.**
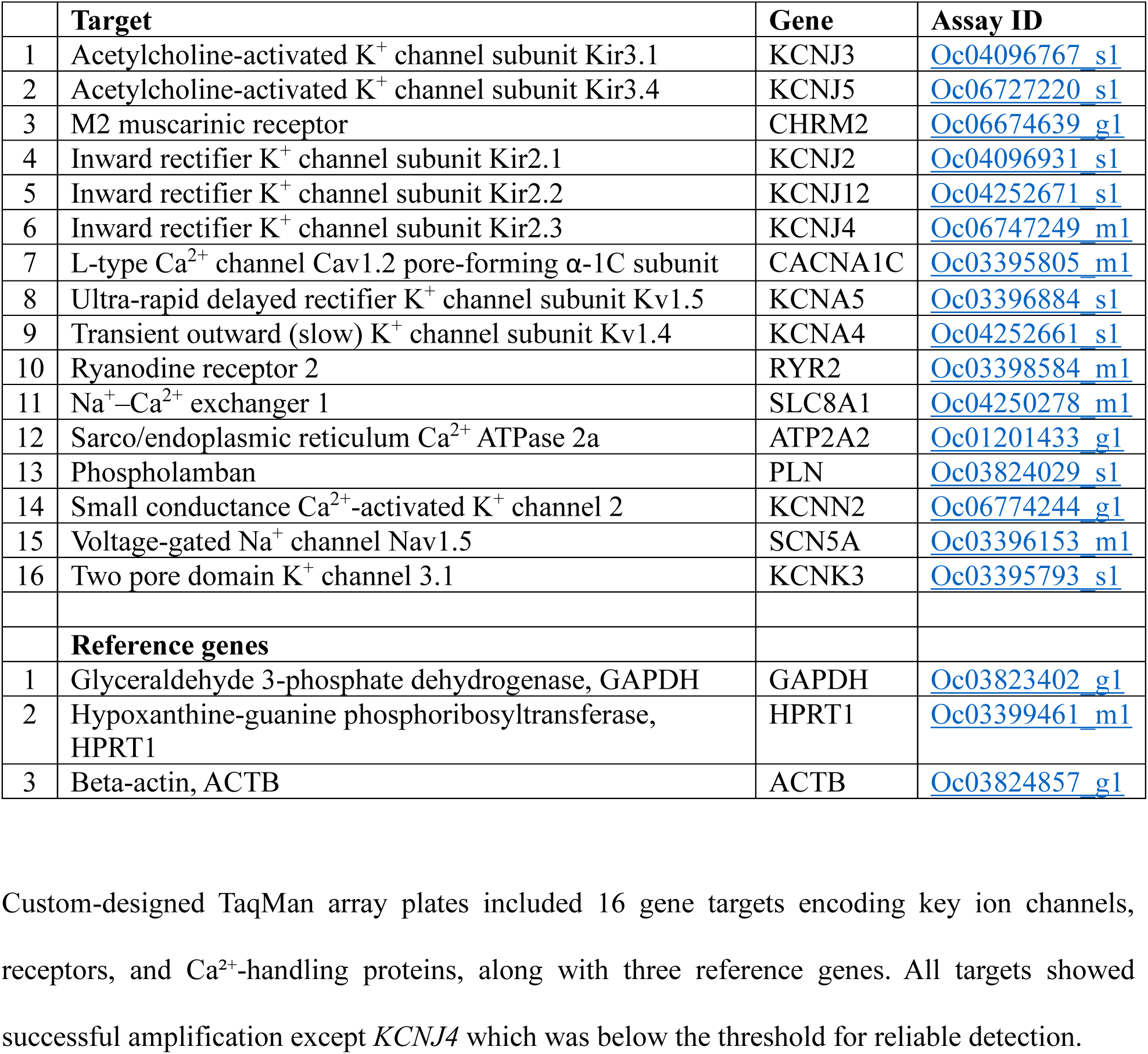
qPCR gene targets.

### Statistics

Statistical analysis was performed using GraphPad Prism 10. Normality was assessed using the Shapiro-Wilk test, and data were transformed when necessary. Statistical significance was evaluated using the appropriate test for each comparison based on data distribution and experimental design as stated in each figure legend, and was attained when *p* < 0.05. Mapping data are presented as mean ± standard error of the mean for *N* animals. qPCR data are presented as median with interquartile range.

## RESULTS

### Baseline atrial electrophysiological properties are comparable between sexes but heterogeneity is greater in females at slower rates

APD_50_ was longer in females compared to males (*p* = 0.046), however no sex differences were observed in ERP, APD_80_, or APD_80_ restitution (Figure 1). Although APD_80_ across the atria was similar between sexes, greater spatial heterogeneity was observed in optical maps from females and regional analysis revealed sex-specific interatrial differences in APD (Figure 2). At a BCL of 300 ms, females exhibited longer APD_50_ and APD_80_ in the RA and IAR compared to males. In both sexes, APD_50_ in the RA and IAR was longer than in the LA; however for APD_80_, regional differences between the LA and RA were only observed in females (Figures 2B and 2C). Although evident at a BCL of 300 ms, these sex differences diminished at faster pacing rates (Figures 2D and 2E). At a BCL of 180 ms, differences in RA and IAR APD between males and females were no longer apparent, and regional differences persisted similarly across sexes (Figures 2F and 2G).

**Figure 2.**
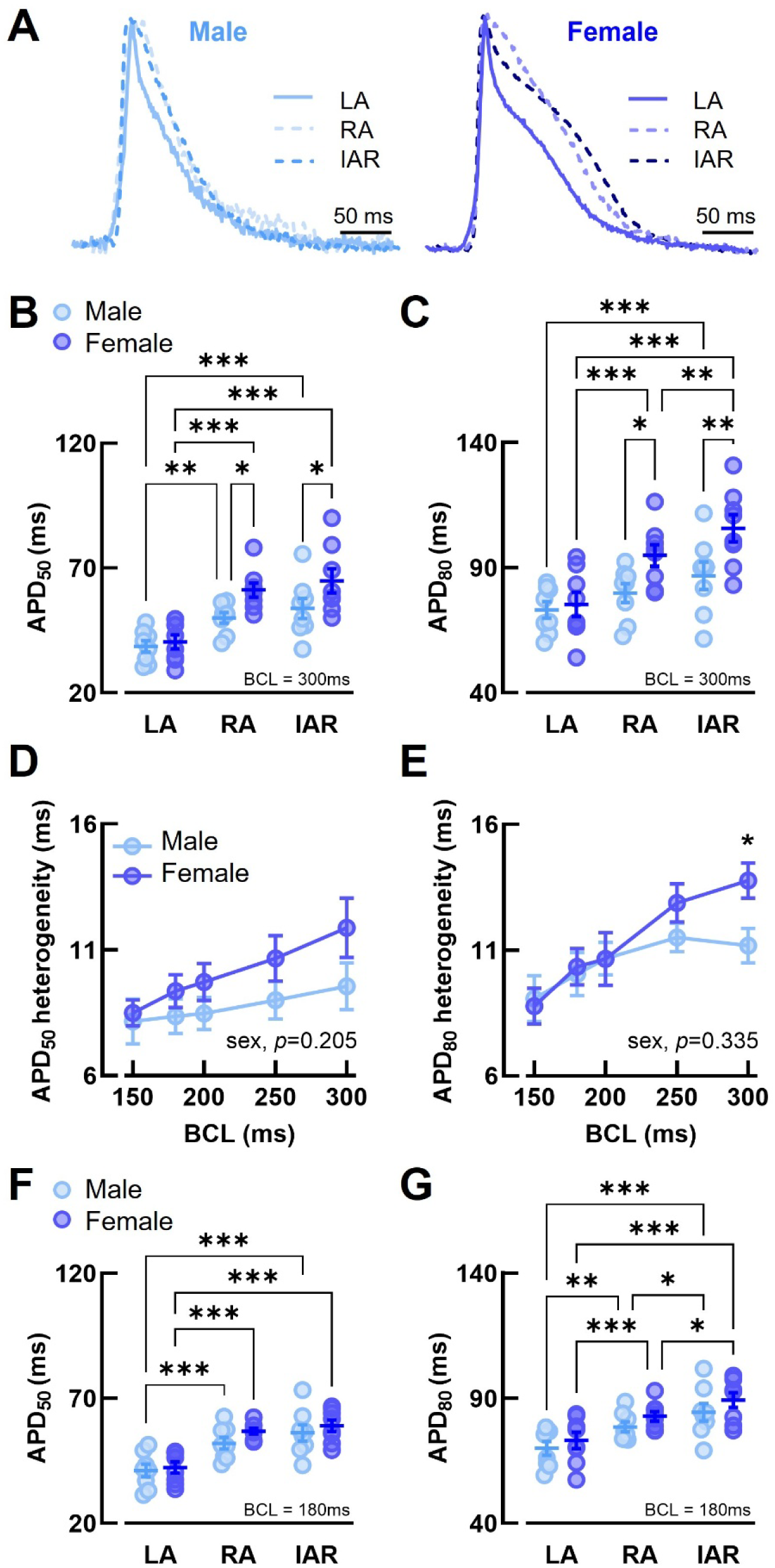
Female rabbit atria have greater APD heterogeneity at slow rates. **(A)** Example action potentials, **(B)** APD_50_ and **(C)** APD_80_ from the LA, RA and IAR of male and female rabbit hearts at BCL = 300 ms. **(D)** APD_50_ heterogeneity and **(E)** APD_80_ heterogeneity across the atria at different BCLs. **(F)** APD_50_ and **(G)** APD_80_ from different atrial regions at BCL = 180 ms. N = 8/group. Statistical significance assessed using two-way ANOVA with repeated measures. APD_50_ = action potential duration at 50% repolarization, APD_80_ = action potential duration at 80% repolarization, BCL = basic cycle length, IAR = interatrial region, LA = left atrium, RA = right atrium.

### Calcium transient duration is prolonged in females

Females had longer CaTD_50_ and CaTD_80_ compared to males (*p* = 0.006 and *p* = 0.005 respectively), along with steeper CaTD_80_ restitution (*p* = 0.021, Figures 3C to 3E). Consistent with longer CaTD and a trend toward prolonged time to peak (*p* = 0.083, Figure 3F), females also showed a tendency for slower recovery of Ca²⁺ release following a premature stimulation (*p* = 0.110, Figures 3G and 3H). No sex differences were observed in CaTD_80_ spatial heterogeneity (Figure 3I), with CaTD longer in females across all atrial regions (Supplemental Figure 1).

**Figure 3.**
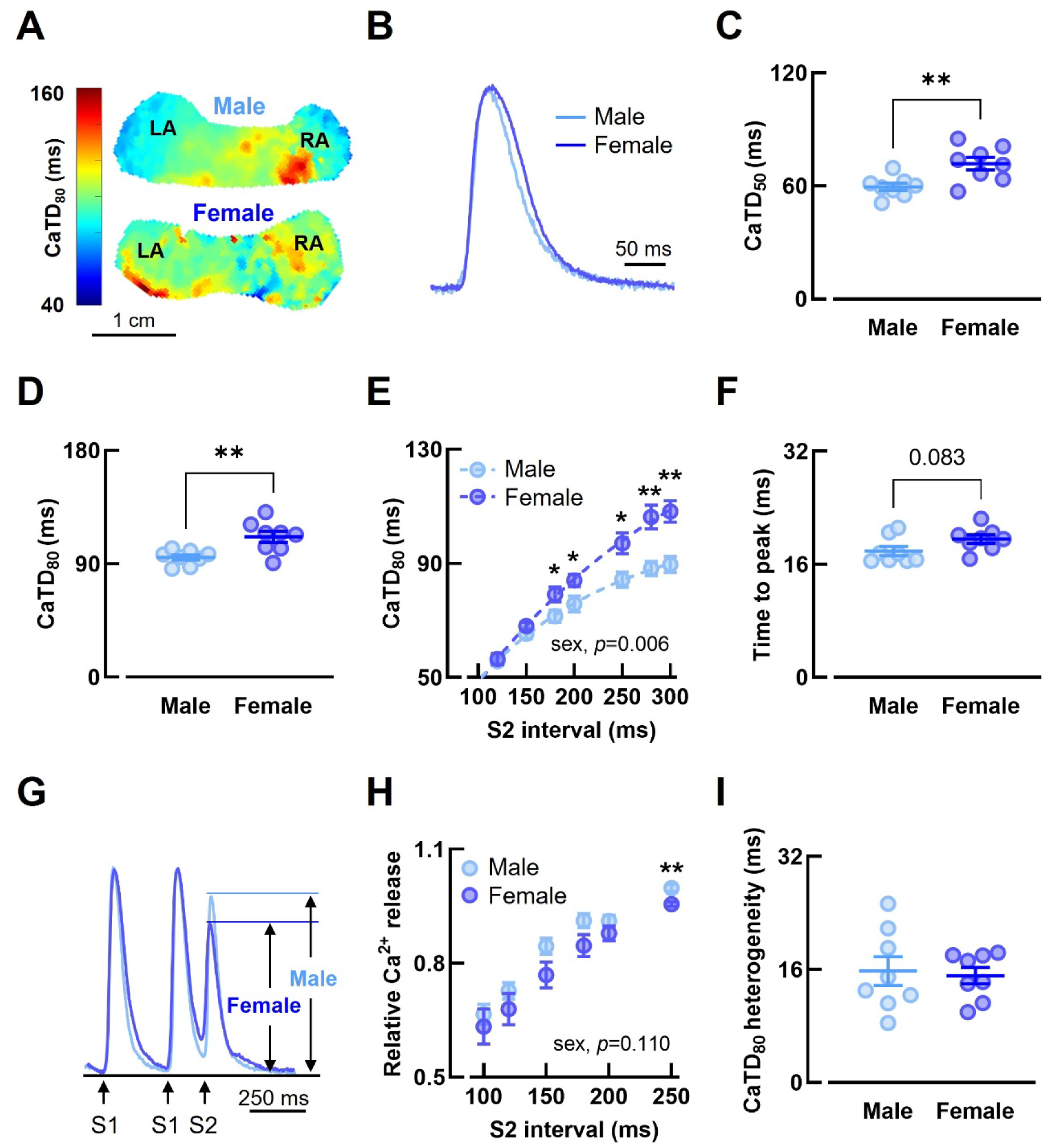
– CaTD is prolonged in female rabbit atria. **(A)** Example CaTD_80_ maps, **(B)** example Ca^2+^ transients, **(C)** CaTD_50_, **(D)** CaTD_80_, **(E)** CaTD_80_ restitution and **(F)** Ca^2+^ transient time to peak in male and female rabbit atria. **(G)** Example Ca^2+^ transient recovery and **(H)** relative Ca^2+^ release following a premature stimulation. **(I)** CaTD_80_ heterogeneity in male and female atria. N = 8/group. Statistical significance assessed using unpaired *t*-test (C, D and I), Mann–Whitney *U* test (F), two-way ANOVA with repeated measures (E) and two-way mixed effects model (H). A, B, C, D, F and I, BCL = 300 ms. E, G and H, S1 interval = 280 ms. BCL = basic cycle length, CaTD_50_ = Ca^2+^ transient duration at 50% decay, CaTD_80_ = Ca^2+^ transient duration at 80% decay.

### Susceptibility to rapid pacing-induced arrhythmias is comparable between sexes, but females are more prone to transient arrhythmias following premature stimulation

In order to assess whether baseline differences in electrophysiological and CaT properties were associated with sex-specific mechanisms of arrhythmia, we evaluated susceptibility to alternans and arrhythmia using rapid pacing, burst pacing with pause, and premature stimulation protocols. In both males and females, CaT alternans emerged before AP alternans, and no differences in alternans threshold were observed between sexes (Supplemental Figure 2). While susceptibility to arrhythmia with burst pacing with pause was similar across sexes (Supplemental Figure 3), females exhibited a higher incidence of transient arrhythmias following premature stimulation (*p* = 0.044, Figure 4B). These transient arrhythmias were more frequently composed of reentry or combined reentry and ectopy (*p* = 0.036), occurred across a number of S2 intervals (*p* = 0.013), and lasted longer in females compared to males (*p* = 0.044, Figures 4C to 4E).

**Figure 4.**
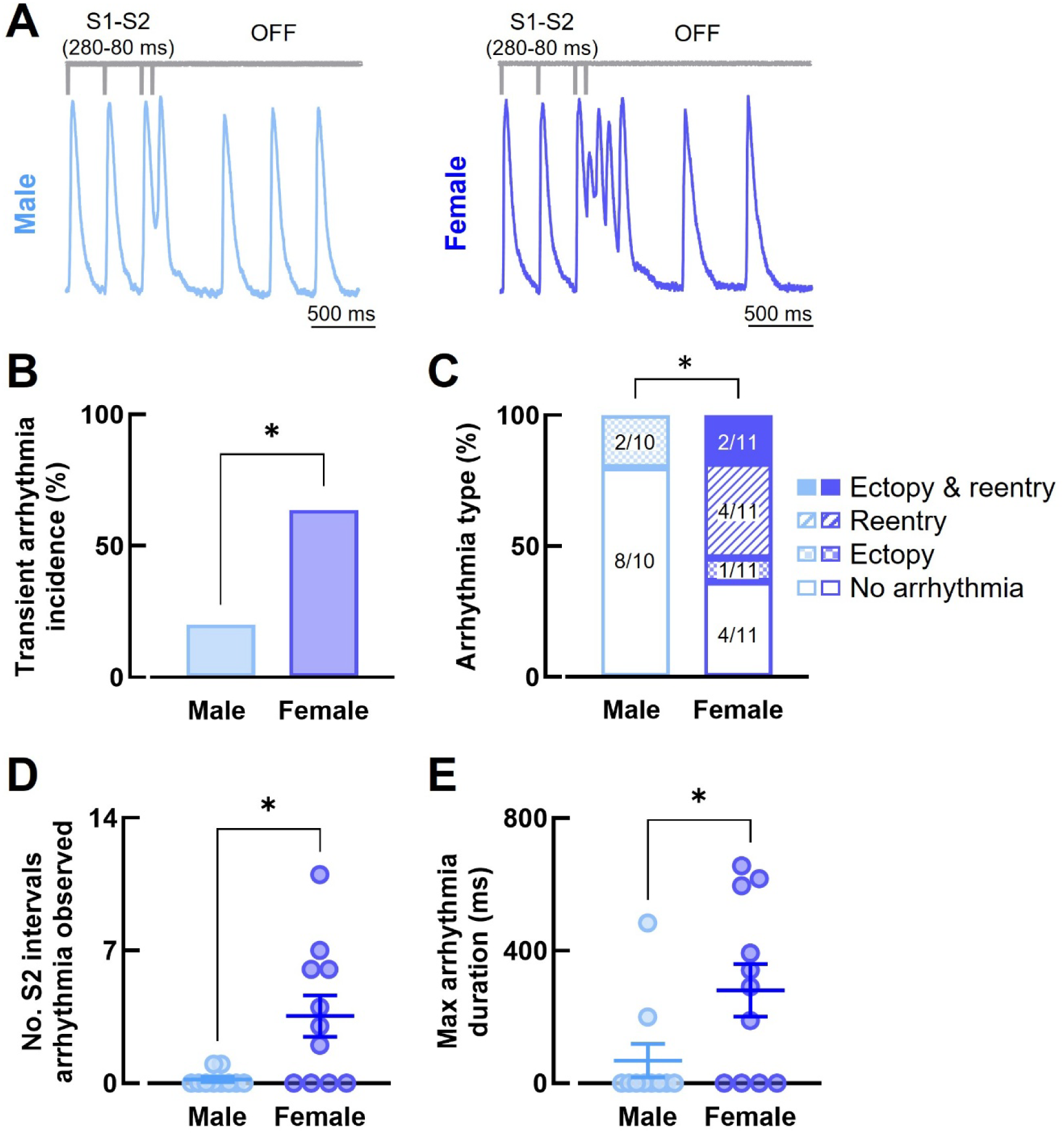
Female rabbit atria are more susceptible to arrhythmia with premature stimulation. **(A)** Example voltage traces, **(B)** incidence of transient arrhythmia, **(C)** arrhythmia type, **(D)** number of S2 intervals arrhythmias were observed, and **(E)** maximum arrhythmia duration during premature stimulation pacing in male and female rabbit atria. N = 10-11/group. Statistical significance assessed using Chi-square test (B), Fisher’s exact test (C) and Mann–Whitney *U* test (D and E). S1 interval = 280 ms.

### Females have attenuated responses to carbachol and less cholinergic-mediated increase in arrhythmia susceptibility than males

To further investigate arrhythmia mechanisms and potential sex differences in responses to parasympathetic modulation, experiments were repeated in the presence of carbachol to assess arrhythmia susceptibility under enhanced vagal tone, a condition known to shorten atrial refractoriness and promote reentrant arrhythmias such as AF.^35^ To assess arrhythmia susceptibility following cholinergic stimulation, burst pacing with pause and premature stimulation protocols were performed as before. Carbachol perfusion increased the incidence and severity of arrhythmias across both pacing protocols (Figure 5). While females exhibited greater susceptibility to transient arrhythmias in response to premature stimulation at baseline, susceptibility increased in males with carbachol, resulting in no sex difference in incidence of transient or sustained arrhythmias under cholinergic conditions using the same protocol (Figure 5B and 5C). In contrast, although no difference was evident between sexes with burst pacing with pause at baseline, males demonstrated a greater tendency toward sustained reentrant arrhythmias following carbachol administration using this protocol (*p* = 0.097, Figure 5E and 5F).

**Figure 5.**
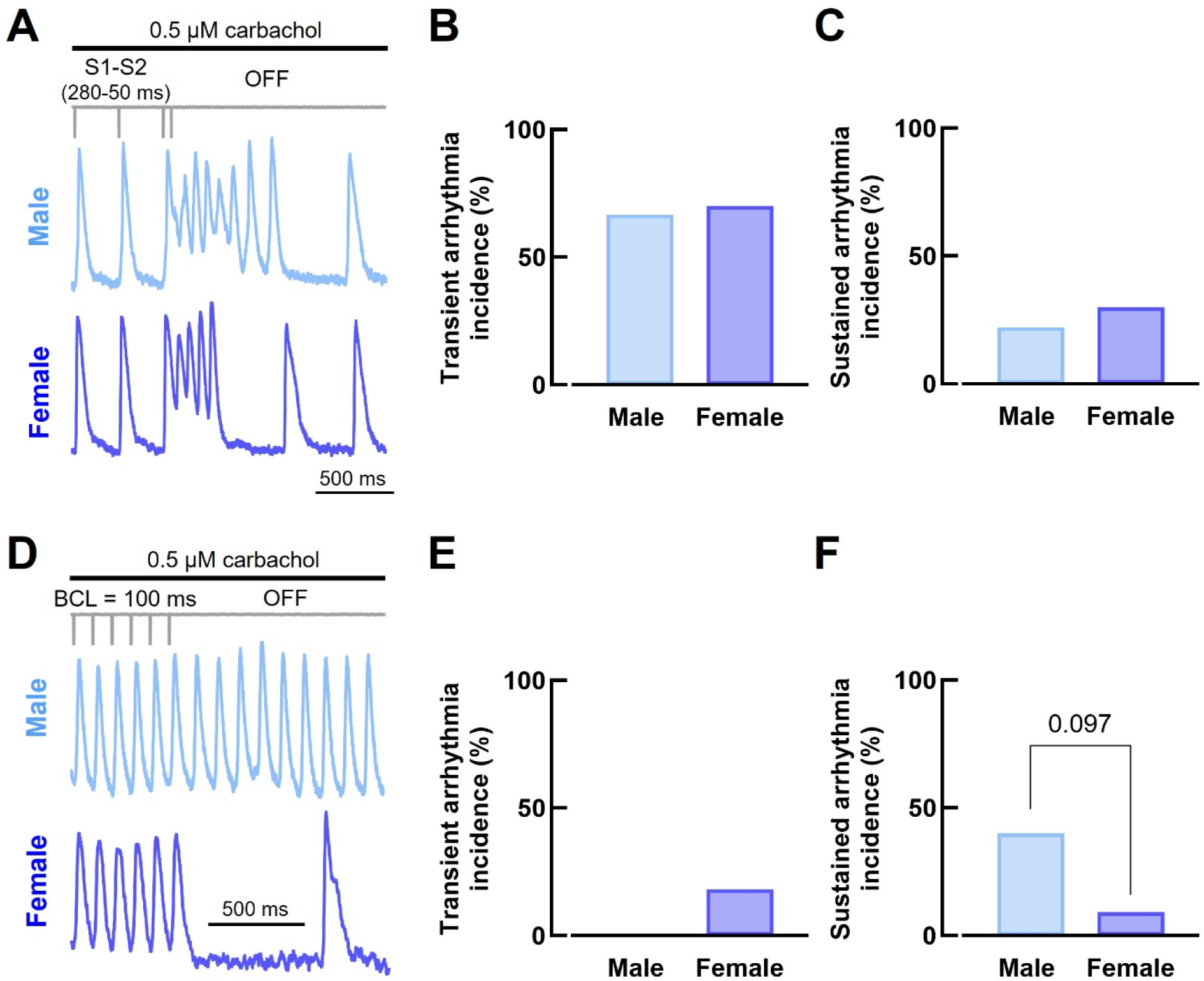
Male rabbit atria have greater tendency for arrhythmia following cholinergic stimulation. **(A)** Example voltage traces, **(B)** incidence of transient and **(C)** sustained arrhythmias during perfusion with carbachol and premature stimulation. **(D)** Example voltage traces, **(E)** incidence of transient and **(F)** sustained arrhythmia during perfusion with carbachol and a burst pacing with pause. A, B and C, N = 9-10/group. S1 interval = 280 ms. D, E and F, N = 10-11/group. Statistical significance assessed using Chi-square test. BCL = 100 and 50 ms.

With carbachol administration, both heart rate and ERP were reduced in males and females (Figures 6A and 6B). While carbachol shortened APD_80_ in males, no change was observed in females (Figure 6D). Additionally, females exhibited longer APD_80_ across a range of S2 intervals following carbachol treatment (Figure 6E). To determine whether these differences were due to regional variations in response to carbachol, we performed regional analysis of the LA, RA and IAR. Post-carbachol, APD_80_ was longer in females compared to males across all atrial regions (Figure 6F). Furthermore, while carbachol shortened APD_80_ in both the RA and LA in males, only the RA showed a response in females (Figures 6G and 6H).

**Figure 6.**
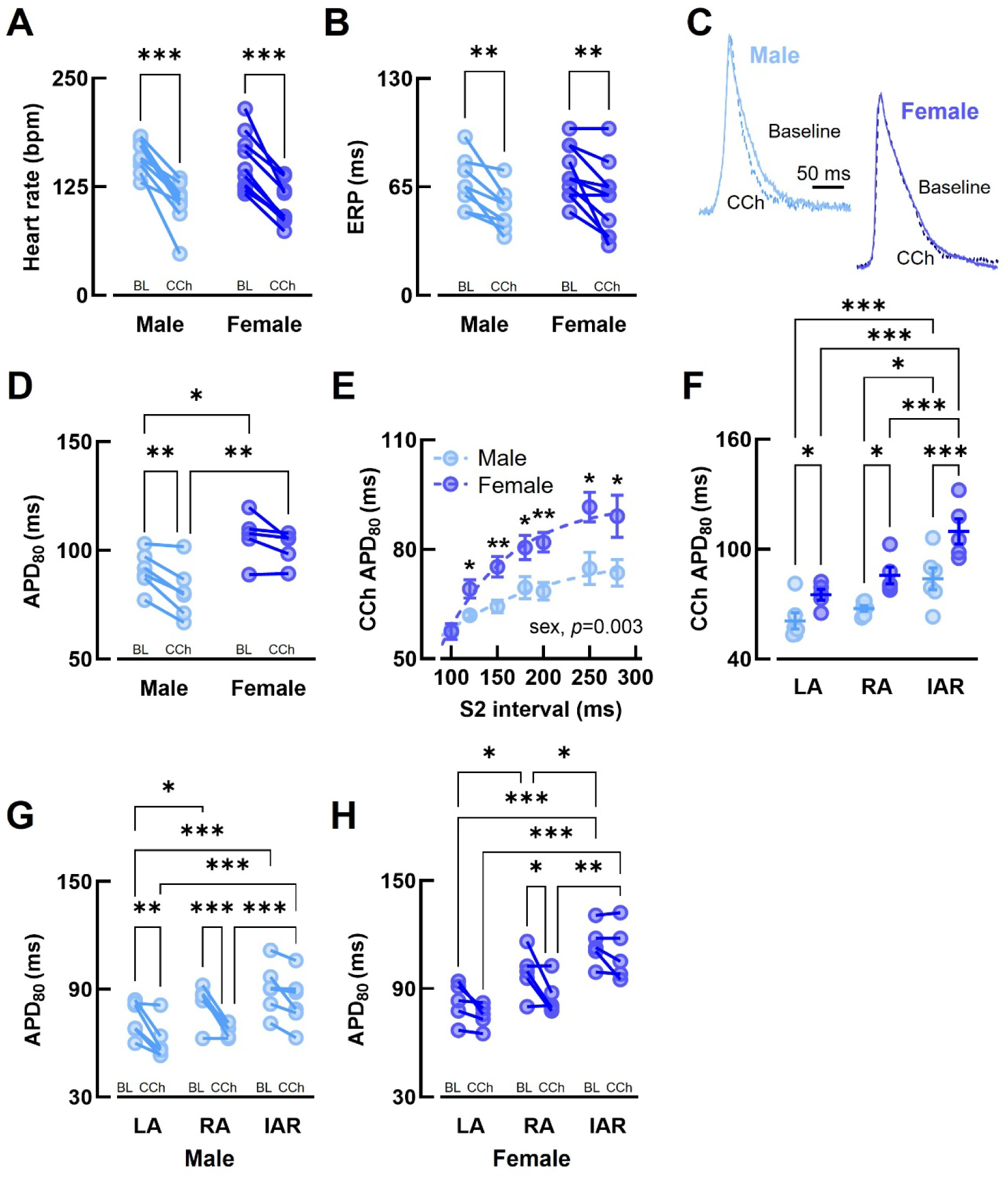
Female rabbit atria have attenuated responses to carbachol. **(A)** Heart rate, **(B)** ERP, **(C)** example action potential traces from the LA, **(D)** APD_80_, and **(E)** APD_80_ restitution in male and female rabbit atria during perfusion with carbachol. **(F)** Comparison of APD_80_ in the LA, RA and IAR of male and female rabbits following carbachol. **(G)** APD_80_ from the LA, RA and IAR of males and **(H)** females at baseline and following carbachol. A and B, N = 9-10/group. C, D, E, F, G, and H, N = 5-6/group. Statistical significance assessed using two-way ANOVA with repeated measures and two-way mixed effects model (E). B and E, S1 interval = 280 ms. C, D, F, G, and H, BCL = 300 ms. APD_80_ = action potential duration at 80% repolarization, BCL = basic cycle length, BL = baseline, bpm = beats per minute, CCh = carbachol, ERP = effective refractory period, IAR = interatrial region, LA = left atrium, RA = right atrium.

### Female atria exhibit greater regional variation in electrophysiology-related gene expression

We next sought to investigate whether sex- and region-specific differences in electrophysiology, CaT properties, and responses to carbachol were associated with underlying variations in gene expression. When atrial chambers were combined and gene expression across the atria was compared by sex, analysis revealed lower expression of *KCNJ12* (inward rectifier K^+^ channel subunit Kir2.2) and *RYR2* (ryanodine receptor 2), with a strong trend toward lower *KCNJ3* (acetylcholine-activated K^+^ channel subunit Kir3.1) expression in females compared to males (Figure 7). Given the greater regional heterogeneity observed in females, we next compared expression in the LA and RA within each sex. In females, *SCN5A* (voltage-gated Na^+^ channel Nav1.5), *RYR2* and *CHRM2* (M2 muscarinic receptor) expression were higher in the RA than in the LA (Figure 8). Additionally, there were trends toward greater expression of *CACNA1C* (L-type Ca^2+^ channel Cav1.2 pore-forming α-1C subunit), *KCNJ12*, *SLC8A1* (Na^+^-Ca^2+^ exchanger), *ATP2A2* (Sarco/endoplasmic reticulum Ca^2+^ ATPase 2a), and *PLN* (phospholamban), along with a trend toward lower *KCNJ2* (inward rectifier K^+^ channel subunit Kir2.1) expression in the RA. In contrast, no significant chamber-specific differences were observed in males, though there were trends toward greater *KCNJ12*, *KCNA5* (ultra-rapid delayed rectifier K^+^ channel subunit Kv1.5), and *KCNJ3* in the RA compared to the LA (Supplemental Figure 4).

**Figure 7.**
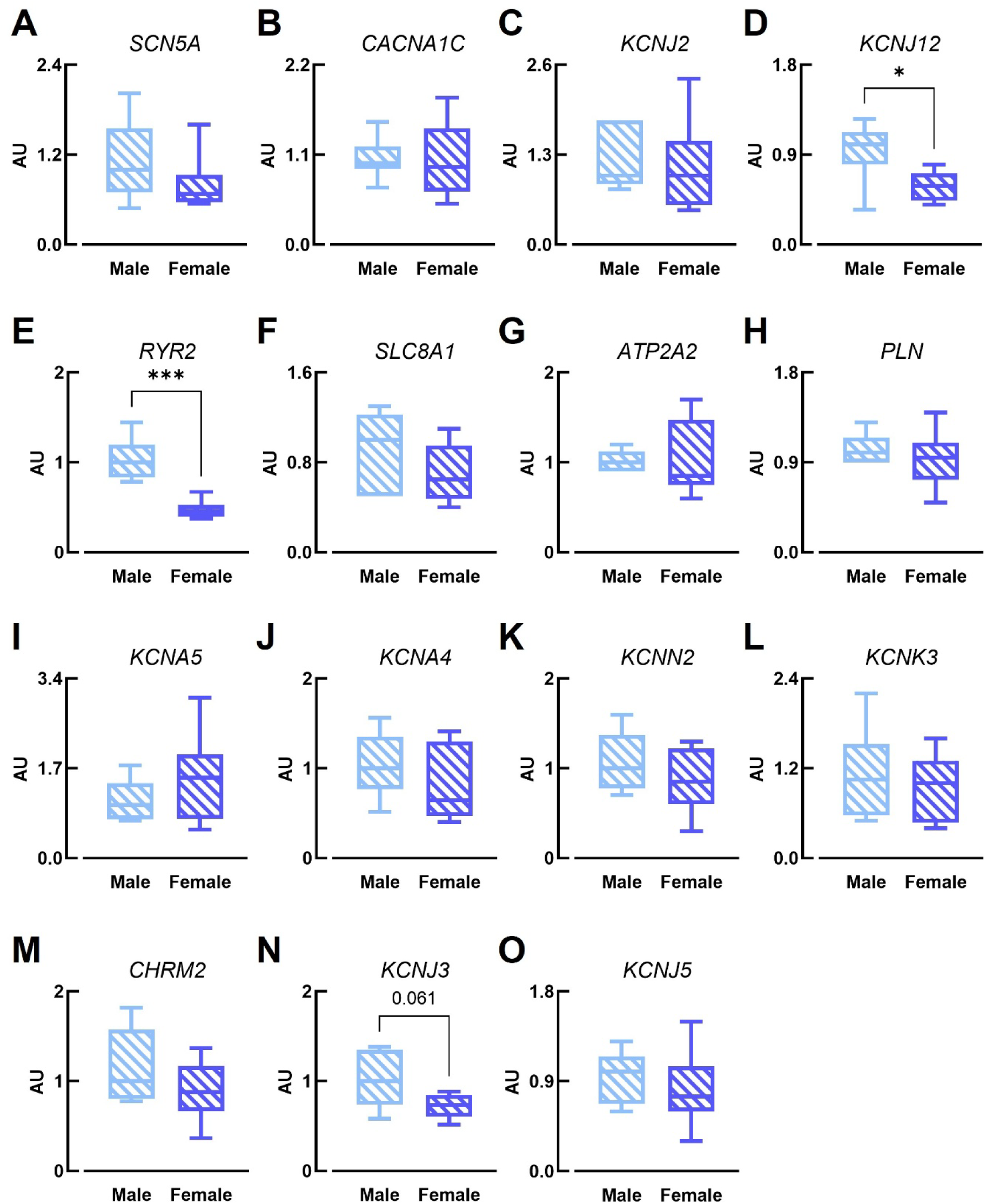
Sex differences in atrial gene expression. **(A)** *SCN5A*, **(B)** *CACNA1C*, **(C)** *KCNJ2*, **(D)** *KCNJ12*, **(E)** *RYR2*, **(F)** *SLC8A1*, **(G)** *ATPA2*, **(H)** *PLN*, **(I)** *KCNA5*, **(J)** *KCNA4*, **(K)** *KCNN2*, **(L)** *KCNK3*, **(M)** *CHRM2*, **(N)** *KCNJ3*, and **(O)** *KCNJ5* expression in male and female atria. N = 6/group performed in triplicate. Statistical significance assessed using Mann–Whitney *U* test (A) and unpaired *t*-test. *SCN5A* = voltage-gated Na^+^ channel Nav1.5, *CACNA1C* = L-type Ca^2+^ channel Cav1.2 pore-forming alpha-1C subunit, *KCNJ2* = inward rectifier K^+^ channel subunit Kir2.1, *KCNJ12* = inward rectifier K^+^ channel subunit Kir2.2, *RYR2* = ryanodine receptor 2, *SLC8A1* = Na^+^–Ca^2+^ exchanger 1, *ATPA2* = sarco/endoplasmic reticulum Ca^2+^ ATPase 2a, *PLN* = phospholamban, *KCNA5* = ultra-rapid delayed rectifier K^+^ channel subunit Kv1.5, *KCNA4* = transient outward (slow) K^+^ channel subunit Kv1.4, *KCNN2* = small conductance Ca^2+^-activated K^+^ channel 2, *KCNK3* = two pore domain K^+^ channel 3.1, *CHRM2* = M2 muscarinic receptor, *KCNJ3* = acetylcholine-activated K^+^ channel subunit Kir3.1, and *KCNJ5* = acetylcholine-activated K^+^ channel subunit Kir3.4.

**Figure 8.**
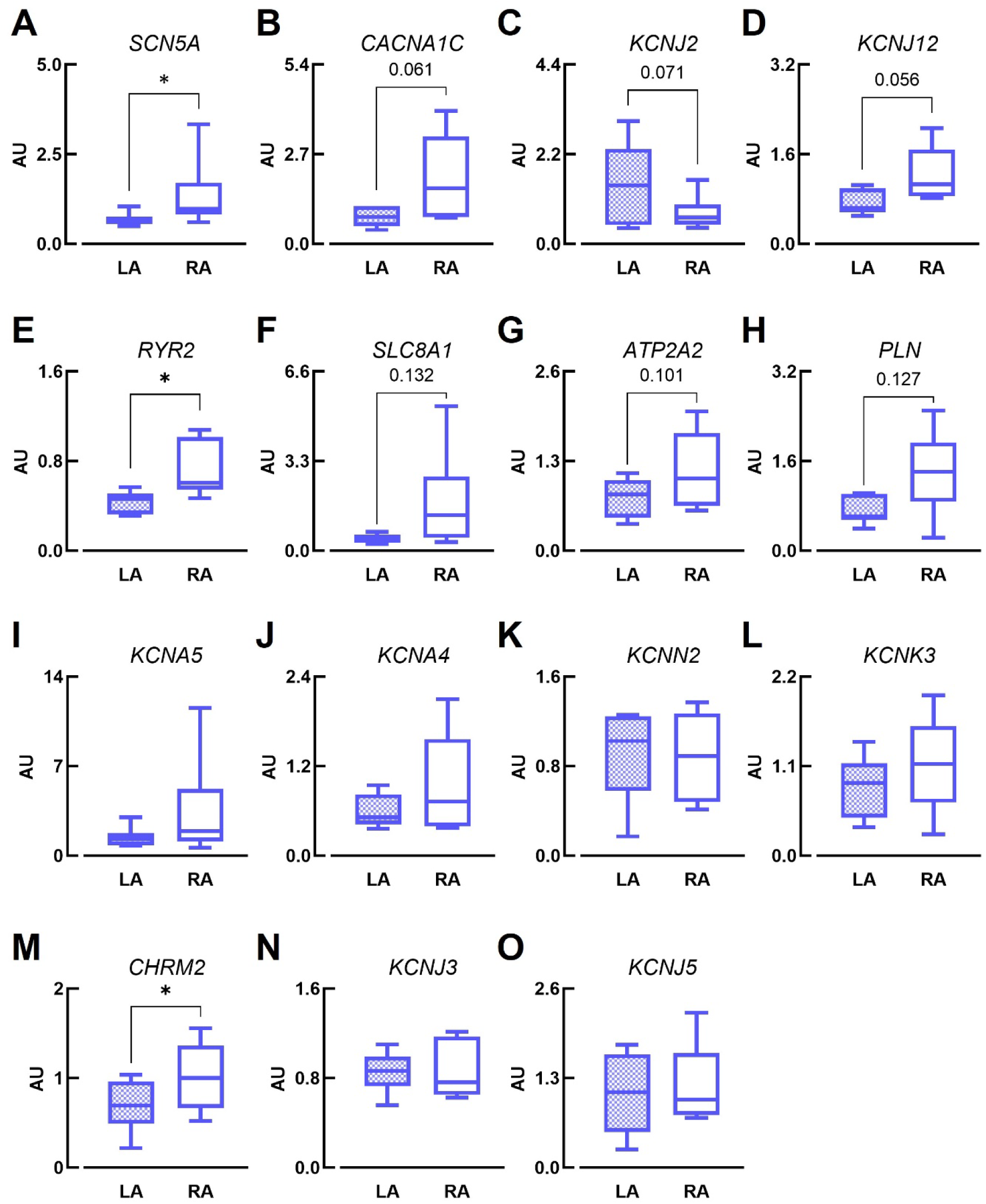
Left vs. right atrial gene expression in females. (A) *SCN5A*, (B) *CACNA1C*, **(C)** *KCNJ2*, **(D)** *KCNJ12*, **(E)** *RYR2*, **(F)** *SLC8A1*, **(G)** *ATPA2*, **(H)** *PLN*, **(I)** *KCNA5*, **(J)** *KCNA4*, **(K)** *KCNN2*, **(L)** *KCNK3*, **(M)** *CHRM2*, **(N)** *KCNJ3*, and **(O)** *KCNJ5* expression in female LA and RA. N = 6/group performed in triplicate. Statistical significance assessed using paired *t*-test. LA = left atrium, RA = right atrium. *SCN5A* = voltage-gated Na^+^ channel Nav1.5, *CACNA1C* = L-type Ca^2+^ channel Cav1.2 pore-forming alpha-1C subunit, *KCNJ2* = inward rectifier K^+^ channel subunit Kir2.1, *KCNJ12* = inward rectifier K^+^ channel subunit Kir2.2, *RYR2* = ryanodine receptor 2, *SLC8A1* = Na^+^–Ca^2+^ exchanger 1, *ATPA2* = sarco/endoplasmic reticulum Ca^2+^ ATPase 2a, *PLN* = phospholamban, *KCNA5* = ultra-rapid delayed rectifier K^+^ channel subunit Kv1.5, *KCNA4* = transient outward (slow) K^+^ channel subunit Kv1.4, *KCNN2* = small conductance Ca^2+^-activated K^+^ channel 2, *KCNK3* = two pore domain K^+^ channel 3.1, *CHRM2* = M2 muscarinic receptor, *KCNJ3* = acetylcholine-activated K^+^ channel subunit Kir3.1, and *KCNJ5* = acetylcholine-activated K^+^ channel subunit Kir3.4.

## DISCUSSION

In this study, we investigated sex differences in AP and CaT properties across the intact atria of a translationally relevant healthy rabbit model. Our findings indicate that female atria exhibit greater APD heterogeneity at slower rates and prolonged CaTD compared to males. Although APD heterogeneity was rate-dependent and comparable to males at shorter BCL, it was associated with greater susceptibility to arrhythmias triggered by premature stimulation at baseline. Interestingly, attenuated responses to carbachol in females, in contrast to the more uniformly stronger responses observed in males, appear protective against cholinergic and burst-pacing-induced arrhythmias.

### Sex differences in APD_50_ and CaTD are independent of gene expression

Consistent with previous studies in mouse, rabbit and human atria^8–11,14,15,36^, we found no sex difference in ERP or APD at late repolarization (e.g. APD_80_), but a longer APD_50_ in females, in agreement with human data.^8,9^ Lower expression of genes encoding K⁺ channel subunits (Kv4.3, KChIP2, Kv1.5, and Kir3.1) may contribute to differences in early repolarization in humans^37^, however besides lower Kir2.2 and tendency for lower Kir3.1 in female rabbits, we found limited sex differences in the expression of K⁺ channel genes across the atria in our study. Similarly, despite the observed prolongation of CaTD across female rabbit atria, the only notable difference in Ca²⁺-handling gene expression was a lower level of *RYR2*. Though lower *RYR2* expression could contribute to prolonged CaTD through less synchronous Ca²⁺ release, reduced release flux, and consequently a smaller driving force for Ca²⁺ reuptake leading to a slower decay^38,39^, it remains unclear whether this alone is sufficient to explain the extent of CaTD prolongation observed.

While gene expression does not necessarily correlate with protein abundance or function, which may itself vary due to membrane insertion, subcellular localization, or post-translational modifications, our results may also reflect sex differences in cellular ultrastructure, as well as channel regulation by hormonal or autonomic influences. Indeed, sex differences have been reported in t-tubule density^40^, autonomic tone^41^, and electrophysiological responses to sex hormones^14^, all of which could additionally contribute to the greater electrophysiological heterogeneity observed in females.

### Greater baseline electrophysiological heterogeneity in females is rate-dependent

While differences in AP morphology between LA and RA rabbit atrial myocytes and greater electrophysiological heterogeneity in the LA of young female mice have previously been reported^14,42^, the rate dependence of interatrial heterogeneity, its underlying mechanisms and implications for arrhythmia susceptibility have not yet been fully characterized. Here, we observed greater interatrial repolarization heterogeneity in females at slower rates, characterized by longer APD in the RA relative to the LA, as well as longer RA and IAR APDs compared with males. These sex differences in APD heterogeneity were diminished at faster rates, where regional differences were similar in both sexes. In females, interatrial electrophysiological differences were associated with differential gene expression between the LA and RA. Importantly, several of the genes upregulated, or showing a strong trend toward upregulation, in female RA versus LA encode ion channels and transporters with rate-dependent properties (e.g., Nav1.5, Cav1.2), providing a potential molecular basis for the observed heterogeneity.

### Greater heterogeneity in female atria at slow rates is associated with enhanced susceptibility to arrhythmia

Heterogeneity in repolarization is typically pro-arrhythmic as it creates regions of delayed repolarization and dispersion of refractoriness that facilitate unidirectional block and reentry. This has been demonstrated in a mouse model of AF where spiral waves preferentially initiated and were sustained at sites with the greatest APD heterogeneity.^43^ While our findings indicate that interatrial heterogeneity in young female rabbits is rate-dependent, suggesting that arrhythmic risk may decrease at faster pacing rates to levels comparable to males, females exhibited greater susceptibility to transient arrhythmias at slower rates with premature stimulation, indicating a wider vulnerable window. Although arrhythmias induced by premature stimuli were short-lived and self-terminating, if comparable delayed afterdepolarization-mediated excitations were to occur in humans, for example, during periods of heightened sympathetic activity, systemic infection, or excessive alcohol intake^44–46^, females may similarly experience an elevated, albeit transient, risk of arrhythmia.

While we have characterized atrial AP morphology, the relationship between electrophysiology and arrhythmia susceptibility is also influenced by structural heterogeneity.^47^ Although females with AF exhibit greater levels of fibrosis^48^, which is associated with slower conduction^49^, collagen deposition is lower in female compared to male RA and comparable between the LA in rabbits of a similar age to those used in our study.^15^ Similarly, larger atrial myocytes and altered connexin localization has been reported in young male mice^10^; however no sex difference in atrial conduction velocity has been observed in either mice or humans.^14,36^ As such, we suggest that the observed differences are more likely attributable to ionic and cellular mechanisms rather than to conduction disturbances secondary to structural differences between sexes in the healthy atria.

### Lower carbachol responsiveness in females attenuates sensitivity to cholinergic-mediated arrhythmias

Susceptibility to and severity of arrhythmias was increased with both premature stimulation and burst pacing protocols following cholinergic activation. Although female rabbit atria demonstrated greater arrhythmia susceptibility following premature stimulation at baseline, males became equally susceptible after carbachol administration and tended to more susceptible to sustained reentry during burst pacing under the same conditions. We suggest that this reflects a greater responsiveness to parasympathetic modulation across the atria in males, as evidenced by a larger degree of carbachol-mediated APD shortening which leads to a shorter wavelength for reentry and facilitates rapid atrial rates and tachyarrhythmia initiation.

Vagal stimulation has previously been shown to induce variable ERP shortening across the atria, thereby increasing dispersion of refractoriness^50^; however, because pacing in this study was performed solely from the RA we cannot directly comment on LA ERP. Nevertheless, since LA APD was not shortened by carbachol in females, the LA likely remained refractory, and in the absence of additional arrhythmic drivers, this may have prevented sustained reentry. In contrast, carbachol shortened APD in both the LA and RA in males, suggesting that, in this context, regional heterogeneity and/or reduced cholinergic responsiveness may be protective in females.

Notably, the two female rabbits that demonstrated transient arrhythmias during burst pacing with pause at baseline also exhibited transient or sustained arrhythmias following carbachol, while baseline data were not collected from one additional female that developed arrhythmia under these conditions. This indicates little change in arrhythmia risk in females, in contrast to males who displayed heightened vulnerability under cholinergic stimulation. Our findings that M2 muscarinic receptor expression is higher in the female RA than LA, and that Kir3.1 expression tends to be lower in females than in males and higher in the male RA than LA, likely underlie these sex-specific differences in carbachol responsiveness. These results align with prior reports of lower Kir3.1 expression in human female atria, larger acetylcholine-activated K⁺ currents, and greater Kir3.1 and 3.4 expression in the human RA compared with the LA.^37,51^ As cholinergic activation has been identified as a key initiator of AF in canine models^52^, these findings raise the possibility that the attenuated cholinergic responses observed in females may limit the formation of a proarrhythmic substrate, thereby acting as a brake on vagally-mediated AF initiation. This is supported by evidence that females typically exhibit higher resting vagal tone, as evidenced by greater high-frequency power of heart rate variability^53^, and by recent findings that female rats require higher-frequency transcutaneous auricular vagus nerve stimulation to decrease heart rate and increase vagally-mediated heart rate variability.^54^ Together, these observations suggest that the female atrium may be intrinsically adapted to tolerate higher parasympathetic drive while limiting arrhythmia susceptibility.

## LIMITATIONS

This study exclusively utilized young adult rabbits, which may not fully capture the spectrum of sex differences that occur across the lifespan. While our findings highlight important sex-based distinctions present in early life, the arrhythmogenic potential in this age group is inherently low, as evidenced by the absence of spontaneous arrhythmias and the inability to induce sustained arrhythmias using pacing protocols, except in the presence of carbachol. Consequently, these results may not be generalizable to aged atria or to other conditions associated with a higher risk of arrhythmia.

Sex hormones are known to play a critical role in modulating cardiac electrophysiology and arrhythmia susceptibility.^14,15^ Although circulating hormone levels were not measured in this study, rabbits are induced ovulators and typically exhibit low, non-cyclical levels of sex hormones under standard laboratory conditions.^15^ This hormonal profile may limit the extent to which hormone-driven sex differences are observed in this model. As a result, it remains unclear whether the differences observed in this study are primarily due to intrinsic biological sex or are influenced by baseline hormonal levels. Future studies incorporating hormonal profiling or experimental manipulation will be important to clarify the specific contributions of sex hormones to electrophysiological variability.

While dual optical mapping enabled assessment of spatial heterogeneity and arrhythmogenic activity in the intact atria, future cellular-level studies are needed to uncover the underlying electrophysiological mechanisms. Similarly, although we performed qPCR on several key gene targets, transcript levels do not necessarily correlate with ion channel or transporter function. Therefore, additional investigations at the protein and functional levels will be essential to fully elucidate the molecular basis of the observed differences.

## CONCLUSIONS

In summary, this study reveals sex- and region-specific differences in atrial electrophysiology, Ca^2+^ dynamics and cholinergic responsiveness in the healthy rabbit heart. Female atria exhibited prolonged CaTD and greater rate-dependent baseline heterogeneity in repolarization, which was associated with greater susceptibility to transient reentrant arrhythmias following premature stimuli. In contrast, males had heightened susceptibility to arrhythmias in the presence of carbachol, consistent with sex- and region-dependent differences in muscarinic receptor and acetylcholine-activated K⁺ channel subunit gene expression. Together, these findings provide new insight into the intrinsic electrophysiological and molecular differences that may underlie sex-specific vulnerability to atrial arrhythmias and underscore the importance of considering sex as a biological variable in cardiac electrophysiology and arrhythmia research.

## ACKNOWLEDGEMENTS

The authors thank Cara Wademan at the UC Davis School of Veterinary Medicine Real-time PCR Research and Diagnostics Core Facility for performing qPCR experiments.

## Abbreviations

AF: atrial fibrillation
AP: action potential
APD: action potential duration
BCL: basic cycle length
CaT: calcium transient
CaTD: calcium transient duration
ERP: effective refractory period
IAR: interatrial region
LA: left atria
RA: right atria

