## Supplemental Material for "Sex-specific electrophysiology and cholinergic responses underlie differential mechanisms of arrhythmia vulnerability in rabbit atria"

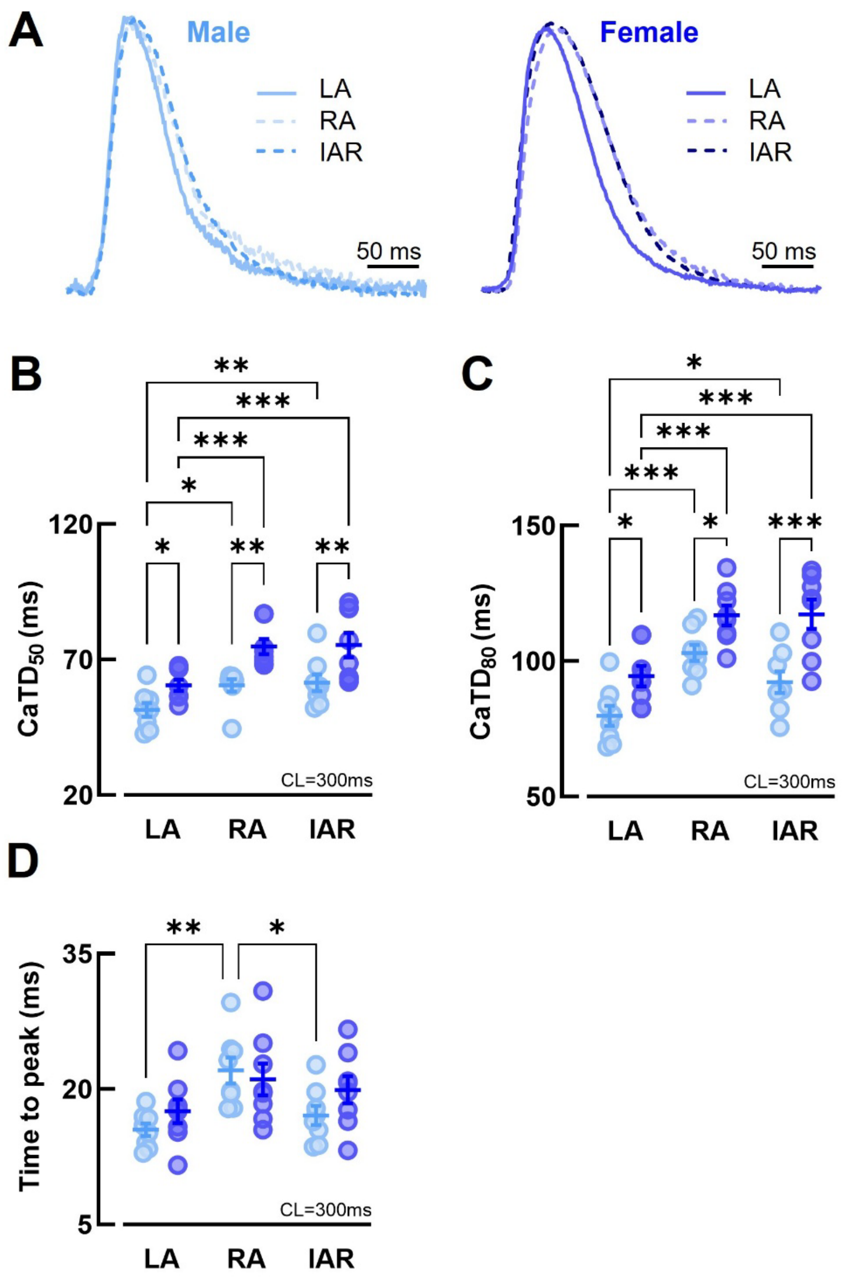


**Supplemental Figure 1 – CaTD is prolonged in females across all atrial regions. (A)** Example $Ca^{2+}$ transients, **(B)** $CaTD_{50}$, **(C)** $CaTD_{80}$ and **(D)** $Ca^{2+}$ transient time to peak from the LA, RA and IAR of male and female rabbits at BCL = 300 ms. N = 8/group. Statistical significance assessed using two-way ANOVA with repeated measures. BCL = basic cycle length, $CaTD_{50}$ = $Ca^{2+}$

transient duration at 50% decay, $CaTD_{80}$ = $Ca^{2+}$ transient duration at 80% decay, IAR = interatrial region, LA = left atrium, RA = right atrium.

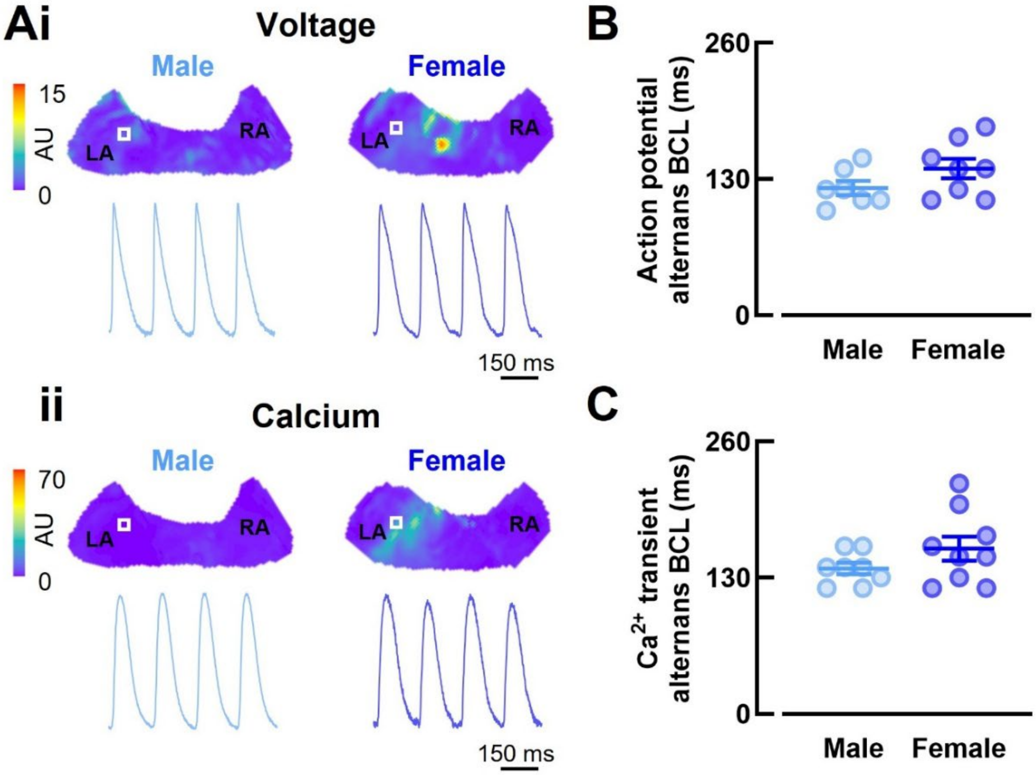


**Supplemental Figure 2 – Alternans susceptibility is similar in male and female rabbit atria.** **(A)** Example voltage (i) and $Ca^{2+}$ (ii) contour maps and corresponding traces taken from a region of interest box in the LA at BCL = 150 ms. **(B)** BCL threshold for action potential alternans and **(C)** BCL threshold for $Ca^{2+}$ transient alternans in male and female atria. N = 8/group. Statistical significance assessed using unpaired *t*-test. BCL = basic cycle length, LA = left atrium, RA = right atrium.

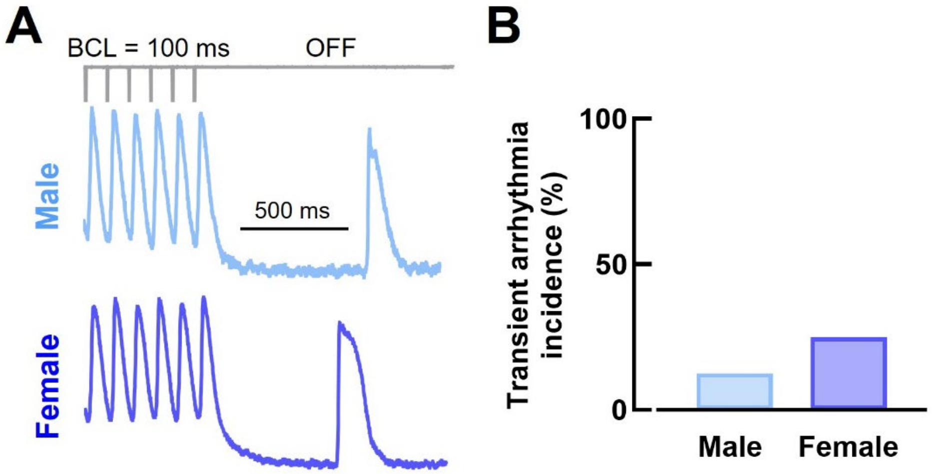


**Supplemental Figure 3 – Susceptibility to arrhythmia with burst pacing with pause is similar between sexes. (A)** Example voltage traces and **(B)** incidence of transient arrhythmia during burst pacing with pause. N = 8/group. Statistical significance assessed using Chi-square test. BCL = 100 and 50 ms. BCL = basic cycle length.

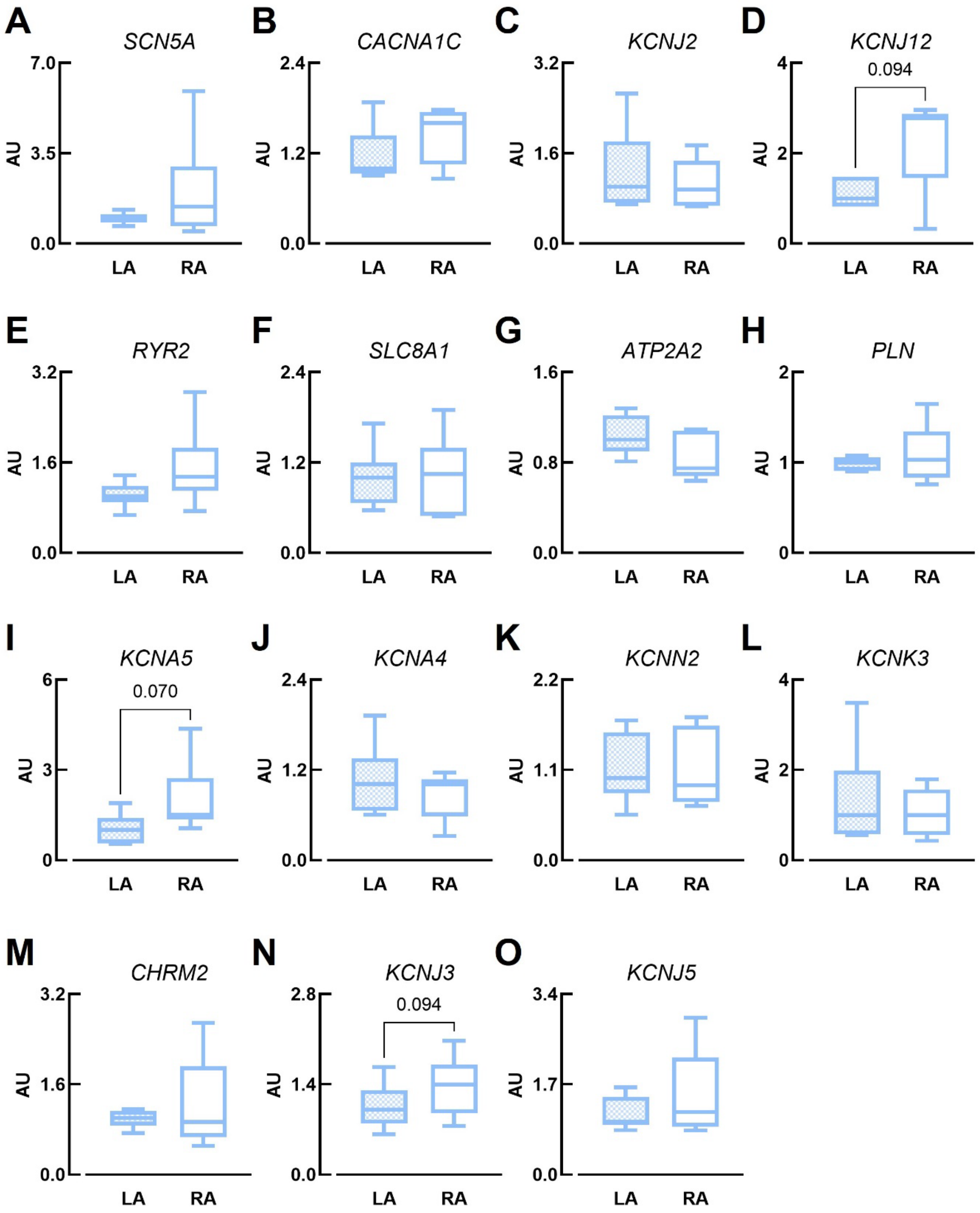


**Supplemental Figure 4 – Left vs. right atrial gene expression in males.** **(A)** *SCN5A*, **(B)** *CACNA1C*, **(C)** *KCNJ2*, **(D)** *KCNJ12*, **(E)** *RYR2*, **(F)** *SLC8A1*, **(G)** *ATPA2*, **(H)** *PLN*, **(I)** *KCNA5*, **(J)** *KCNA4*, **(K)** *KCNN2*, **(L)** *KCNK3*, **(M)** *CHRM2*, **(N)** *KCNJ3*, and **(O)** *KCNJ5* expression in male LA and RA. N = 6/group performed in triplicate. Statistical significance assessed using paired *t*-test and Wilcoxon matched-pairs signed rank test (D). LA = left atrium, RA = right atrium. *SCN5A* = voltage-gated $Na^+$ channel Nav1.5, *CACNA1C* = L-type $Ca^{2+}$ channel Cav1.2 pore-forming alpha-1C subunit, *KCNJ2* = inward rectifier $K^+$ channel subunit Kir2.1, *KCNJ12* = inward rectifier $K^+$ channel subunit Kir2.2, *RYR2* = ryanodine receptor 2, *SLC8A1* = $Na^+$–$Ca^{2+}$ exchanger 1, *ATPA2* = sarco/endoplasmic reticulum $Ca^{2+}$ ATPase 2a, *PLN* = phospholamban, *KCNA5* = ultra-rapid delayed rectifier $K^+$ channel subunit Kv1.5, *KCNA4* = transient outward (slow) $K^+$ channel subunit Kv1.4, *KCNN2* = small conductance $Ca^{2+}$-activated $K^+$ channel 2, *KCNK3* = two pore domain $K^+$ channel 3.1, *CHRM2* = M2 muscarinic receptor, *KCNJ3* = acetylcholine-activated $K^+$ channel subunit Kir3.1, and *KCNJ5* = acetylcholine-activated $K^+$ channel subunit Kir3.4.
